# Anatomy-preserving rare-subregion correction for pediatric brain tumor segmentation on multiparametric MRI: a BraTS-PEDs 2026 challenge study

**DOI:** 10.64898/2026.11.11.765920

**Authors:** Hari Mohan Rai, Dauren Omarbekov, Maria Gul

## Abstract

Pediatric brain tumor subregion segmentation is strongly imbalanced: whole-tumor anatomy is comparatively stable, whereas enhancing tumor (ET), cystic component (CC), and edema (ED) may be absent or small. We present an anatomy-preserving multi-stage system for BraTS-PEDs 2026 Task 2. A five-fold residual-encoder nnU-Net supplies robust anatomy and calibrated out-of-fold priors. Candidate ET and CC regions are recovered by independent fold agreement and component-level filters. A rare-aware residual adapter then operates only inside voxels labeled non-enhancing tumor core (NETC), using MRI, signed intermodality contrasts, probability uncertainty, spatial anatomy, and coordinates. Its constrained assembly permits only NETC-to-ET or NETC-to-CC transitions; therefore background, ED, tumor core, and whole tumor are invariant. On 236 out-of-fold cases, the adapter improved CC Dice from 0.223 to 0.253 and CC normalized surface Dice from 0.233 to 0.263, while preserving tumor-core and whole-tumor scores exactly. The frozen Docker submission achieved Dice scores of 0.510, 0.908, 0.144, 0.000, 0.937, and 0.937 for ET, NETC, CC, ED, tumor core, and whole tumor, respectively, on the 91-case challenge validation set. The results show that constrained rare-subtype correction can improve difficult labels without sacrificing strong anatomical predictions, while also exposing persistent domain shift for CC and ED.

## 1 Introduction

Accurate delineation of pediatric brain tumor subregions is difficult because the target classes differ greatly in prevalence, volume, appearance, and clinical meaning. A conventional flat multiclass objective is dominated by background and large tumor regions. In our labeled cohort, ET was present in 160/236 cases at a 10-voxel threshold, while CC and ED were present in only 76/236 and 50/236 cases. Consequently, a model can obtain strong tumor-core (TC) and whole-tumor (WT) scores while suppressing genuine rare tissue or hallucinating it in absent cases.

nnU-Net provides a strong self-configuring baseline for biomedical segmentation [1], building on the encoder–decoder U-Net architecture [6], and remains highly effective for gross tumor anatomy. Our error analysis, however, showed that many ET and CC errors were not failures to identify tumor. Instead, rare-subtype voxels were absorbed into NETC. This motivated a narrower task: preserve the reliable anatomy and learn only whether selected NETC voxels should be promoted to ET or CC.

Our contribution is a deployable, anatomy-preserving pipeline with three ideas: (i) true out-of-fold (OOF) nnU-Net priors for unbiased error learning; (ii) component consensus that recovers supported rare candidates; and (iii) a residual 3D adapter whose output space is explicitly restricted to NETC *→* ET/CC. The restriction makes TC and WT invariant by construction, rather than relying on a soft regularizer. Figure 1 summarizes the frozen Docker system.

**Fig. 1.**
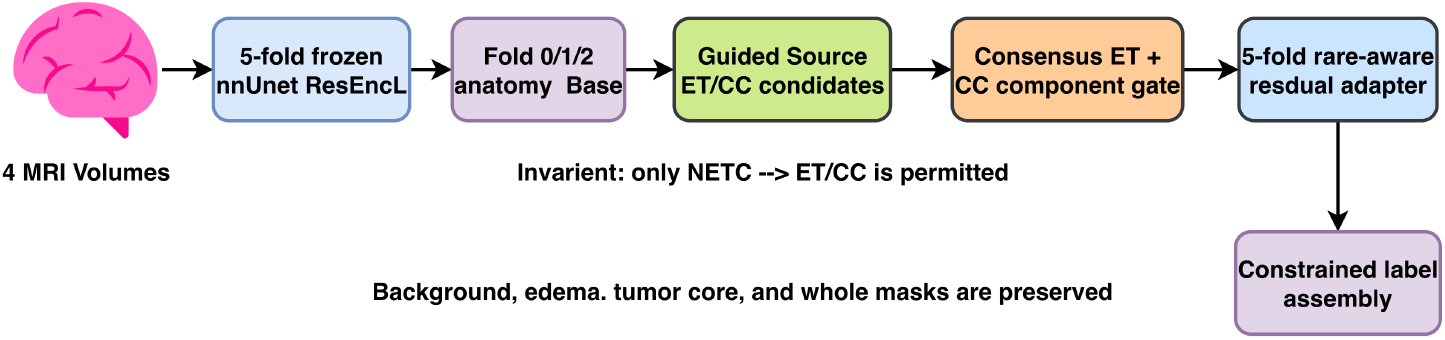
Frozen Docker inference pipeline. The final stages can only relabel NETC as ET or CC. Background, ED, TC, and WT are preserved exactly.

## 2 Materials and Methods

### 2.1 Data and Label Contract

We used 236 labeled training cases, a disjoint 58-case internal holdout for development, and 91 unlabeled challenge validation cases. Each case contained T1-weighted native (T1N), contrast-enhanced T1-weighted (T1C), T2-weighted (T2W), and T2-FLAIR (T2F) MRI [4, 3]. In accordance with the challenge rules, we also cite the MedPerf federated benchmarking framework [2]. No challenge validation labels were used for training.

The official integer labels were ET=1, NETC=2, CC=3, and ED=4. Composite masks were

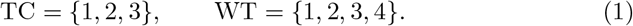

This contract was enforced by executable tests before training and packaging. The phenotype distribution is shown in Fig. 2.

**Fig. 2.**
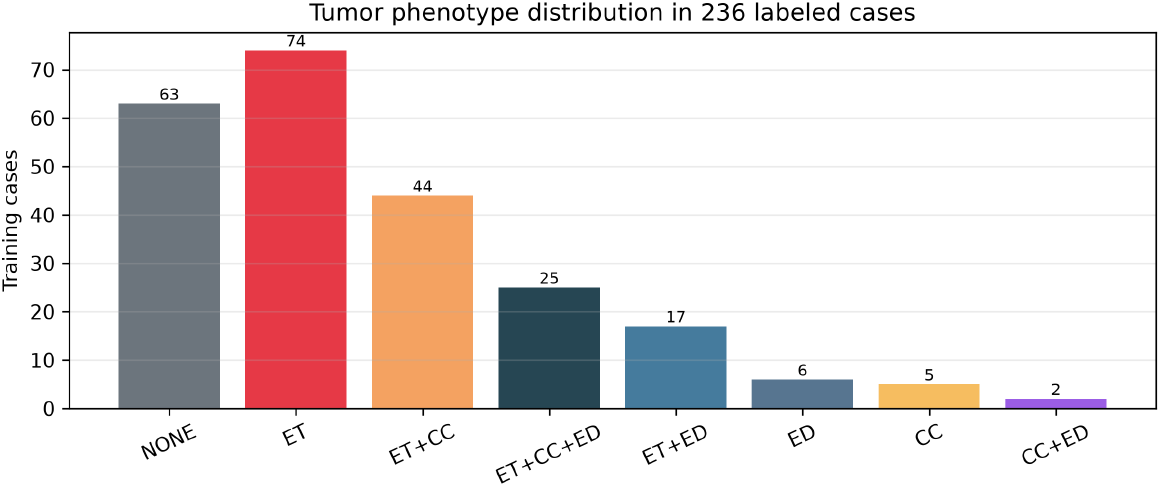
Training phenotype distribution. CC and ED phenotypes are substantially rarer than ET-only and rare-subtype-absent cases.

### 2.2 Frozen Anatomical Teacher

We trained a five-fold 3D full-resolution nnU-Net with a residual-encoder large configuration. Each fold generated OOF predictions and class probabilities only for cases excluded from its training subset. These predictions supplied unbiased failure patterns for adapter training. At Docker inference, folds 0–2 were ensembled directly to form the anatomical base, while all five fold probabilities were available to the learned adapter. The base retained the favorable bias of nnU-Net for TC and WT.

### 2.3 Signed Contrast Features

We computed two signed modality differences,

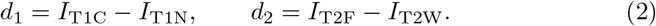

For each difference volume, positive and negative values were normalized separately:

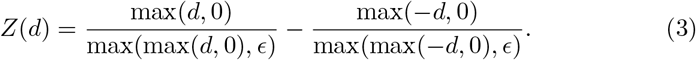

Equation 3 preserves sign and fixes zero as the boundary between enhancement and suppression. Intensity shift and additive noise were disabled for these derived channels because they move that zero boundary. Signed ratios and local contrast channels complemented *Z*(*d*); ZSplit was treated as an auxiliary cue, not as an independent segmentation solution.

### 2.4 Candidate Recovery Before the Adapter

A guided 3D refiner generated candidate ET and CC components from 16 channels: four MRI channels, two ZSplit channels, seven base-prior channels, and three coordinates. Its 160^3^ ROI was centered on the base tumor.

For ET, each candidate connected component *C* was accepted when either fold 1 or fold 2 supported at least 10% of its voxels:

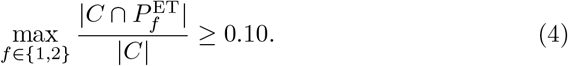

Accepted voxels could modify only base NETC. For CC, a source component required at least 500 promotable NETC voxels and a promotable fraction of at least 0.25. These rules were selected on labeled development data before challenge inference.

### 2.5 Rare-Aware Residual Adapter

The adapter received a 160^3^ ROI with 32 channels (Table 1). A residual 3D encoder–decoder with base width 24 produced ET and CC promotion logits, boundary logits, presence logits, and deep-supervision outputs.

**Table 1.** Adapter input groups.

| Group | Channels | Contents |
|---|---|---|
| MRI | 4 | normalized T1N, T1C, T2W, T2F |
| Signed contrast | 6 | two ZSplit, two signed ratios, two local contrasts |
| Base priors | 7 | official-label and composite nnU-Net priors |
| Probability | 9 | five class probabilities, rare-vs-NETC margins, entropy |
| Anatomy | 3 | TC/NETC interior distance and TC boundary |
| Position | 3 | normalized *x*, *y*, *z* coordinates |
| Total | 32 |  |

The promotion target for subtype *r ∈ {* ET, CC*}* was positive only where the OOF base predicted NETC and the reference label was *r*. The objective combined focal loss [5], Tversky loss [7], generalized Dice loss [8], boundary focal loss, case-level presence loss, identity loss, a conversion-budget penalty, and deep supervision:

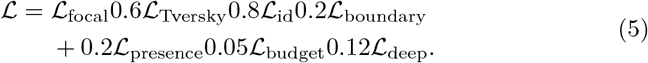

The Tversky parameters were *α* = 0.30 and *β* = 0.75, emphasizing false negatives. ET/CC promotion positive weights were 5/10 and presence positive weights were 0.5/2.4. Training used AdamW, batch size one, mixed precision, a cosine learning-rate schedule, and metric-based checkpoint audits.

For each fold, three specialist checkpoints selected from OOF ET or CC behavior were averaged. Across folds, probabilities were averaged only where that fold considered the voxel eligible NETC. ET used promotion/presence thresholds 0.70/0.50 and CC used 0.50/0.80.

#### Algorithm 1

Docker inference for one case

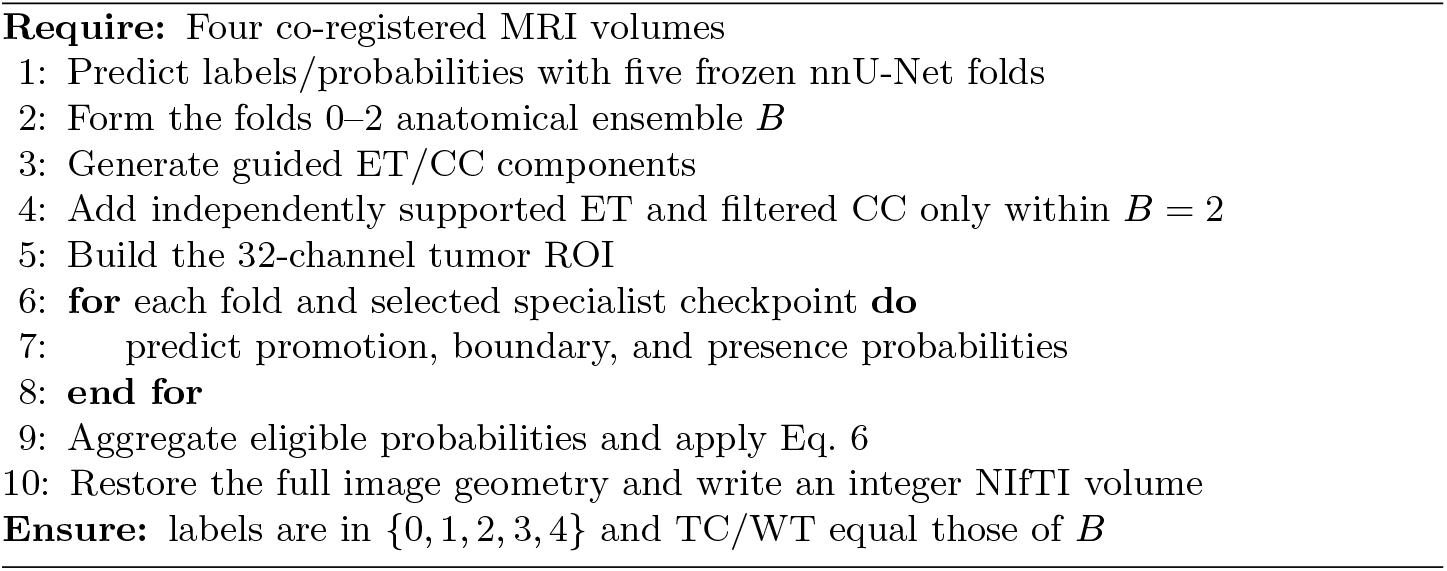

### 2.6 Constrained Assembly

Let *B*(*x*) be the base label and 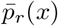 and 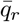 be the aggregated voxel and presence probabilities. The final segmentation is

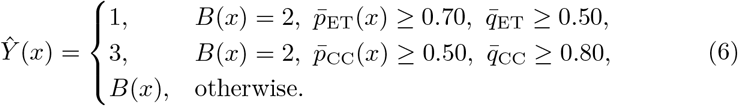

Conflicts were resolved by calibrated rare-class confidence. Since every allowed transition remains inside {1, 2, 3}, Eq. 6 guarantees exact TC and WT preservation and cannot create ED.

### 2.7 Evaluation

We report lesion-wise instance F1, global Dice similarity coefficient (DSC), normalized surface Dice (NSD), and 95th-percentile Hausdorff distance (HD95). OOF evaluation pooled all 236 training cases, with every prediction generated by a model that had not trained on that case. The challenge validation numbers were returned by the official server. The Docker container was additionally tested offline with networking disabled, read-only input and root filesystems, and a single GPU.

## 3 Results

### 3.1 Out-of-Fold Ablation

Table 2 and Fig. 3 compare the five-fold nnU-Net base with the rare-aware adapter. CC improved consistently across DSC, NSD, instance F1, and HD95. ET improved modestly. NETC decreased slightly because promoted voxels are removed from NETC. ED, TC, and WT were exactly unchanged.

**Table 2.** OOF results on 236 labeled cases. HD95 is in mm.

| Metric | Method | ET | NETC | CC | ED | TC | WT |
|---|---|---|---|---|---|---|---|
| Instance F1 | nnU-Net | .502 | .718 | .184 | .129 | .770 | .781 |
|  | + adapter | .500 | .706 | **.215** | .129 | .770 | .781 |
| DSC | nnU-Net | .549 | .769 | .223 | .177 | .846 | .864 |
|  | + adapter | **.553** | .767 | **.253** | .177 | .846 | .864 |
| NSD | nnU-Net | .567 | .679 | .233 | .147 | .692 | .719 |
|  | + adapter | **.570** | .675 | **.263** | .147 | .692 | .719 |
| HD95 | nnU-Net | 88.98 | 13.52 | 208.82 | 245.32 | 10.12 | 8.38 |
|  | + adapter | **86.86** | 13.52 | **192.51** | 245.32 | 10.12 | 8.38 |

**Fig. 3.**
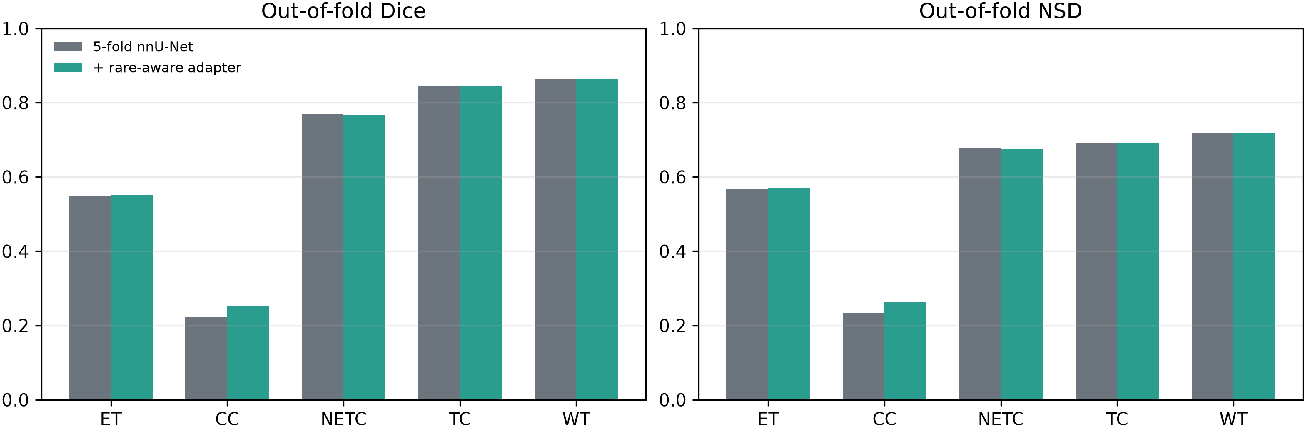
OOF DSC and NSD before and after rare-aware adaptation. TC and WT are unchanged by construction.

### 3.2 Challenge Validation

The frozen Docker candidate’s official 91-case results are shown in Table 3. TC, WT, and NETC remained strong. ET was moderate, CC remained difficult, and ED was not recovered by the constrained ET/CC adapter.

**Table 3.** Official challenge validation results for the frozen Docker candidate.

| Metric | ET | NETC | CC | ED | TC | WT |
|---|---|---|---|---|---|---|
| Instance F1 | .464 | .848 | .198 | .000 | .926 | .926 |
| DSC | .510 | .908 | .144 | .000 | .937 | .937 |
| NSD | .386 | .594 | .107 | .000 | .615 | .615 |
| HD95 (mm) | 123.25 | 7.74 | 262.98 | 373.00 | 7.15 | 7.14 |

**Table 4.**
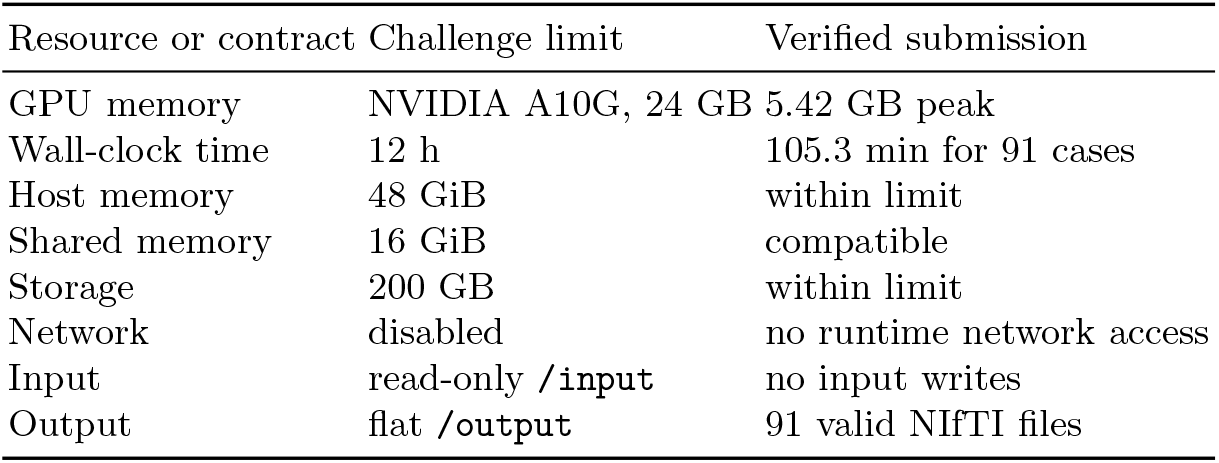
Container compliance against the Task 2 evaluation limits.

| Resource or contract | Challenge limit | Verified submission |
|---|---|---|
| GPU memory | NVIDIA A10G, 24 GB | 5.42 GB peak |
| Wall-clock time | 12 h | 105.3 min for 91 cases |
| Host memory | 48 GiB | within limit |
| Shared memory | 16 GiB | compatible |
| Storage | 200 GB | within limit |
| Network | disabled | no runtime network access |
| Input | read-only /input | no input writes |
| Output | flat /output | 91 valid NIfTI files |

### 3.3 Qualitative Analysis and Deployment

Figure 4 presents OOF examples selected by per-case metric change: a strong ET recovery, a strong CC recovery, and a failure case. The adapter generally converts localized NETC regions without changing gross tumor extent; failures arise when the MRI appearance and fold probabilities provide insufficient separation between CC and NETC.

**Fig. 4.**
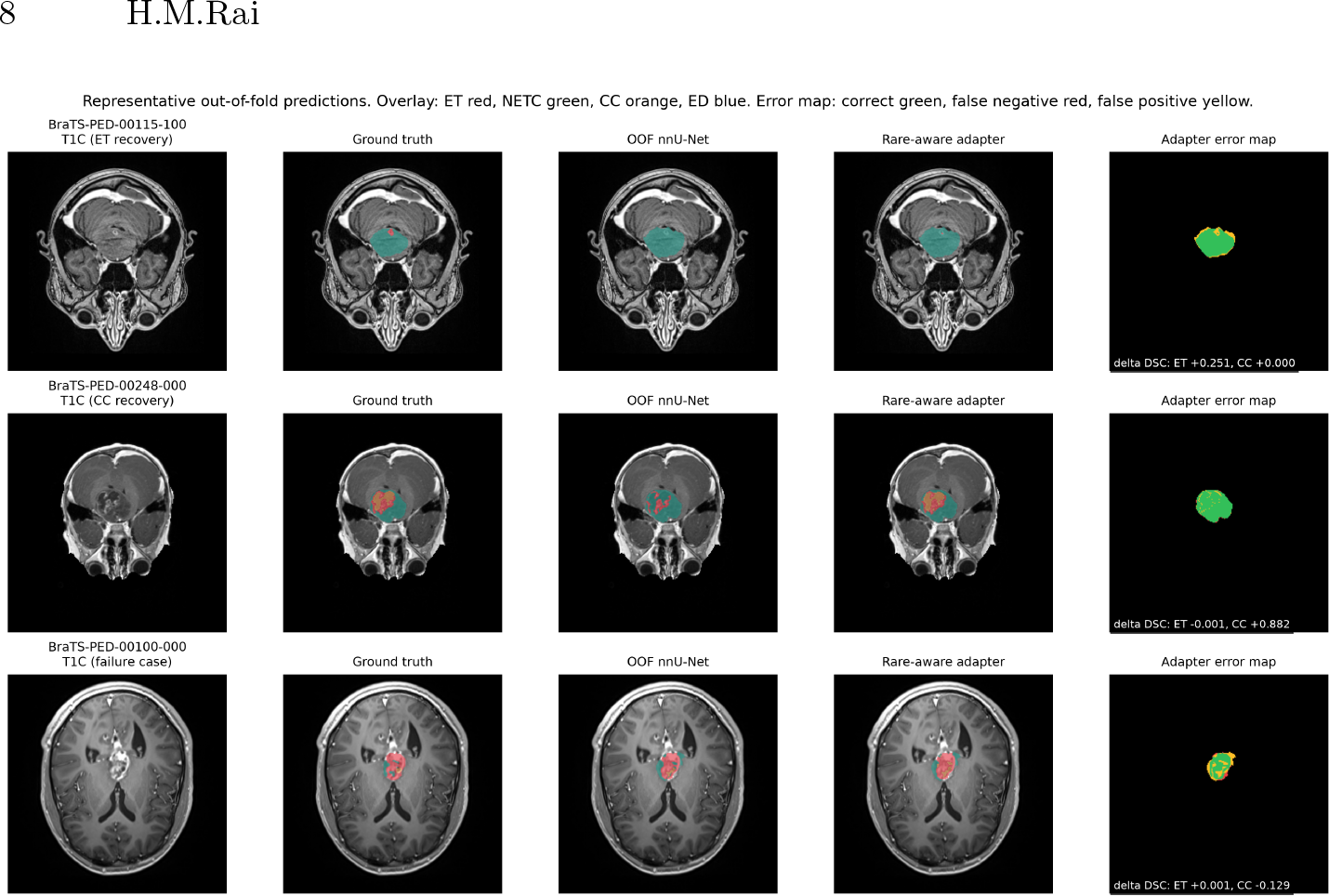
Representative OOF predictions, including two improvements and one failure case. Green error-map pixels are correct, red are false negatives, and yellow are false positives.

The final container completed all 91 validation cases in 105.3 minutes on one GPU. Peak measured CUDA memory during adapter inference was 5.42 GB. Offline, network-disabled execution, read-only filesystem behavior, output count, geometry, affine, datatype, and label-contract checks all passed.

## 4 Discussion

The central result is not that a second network universally outperformed nnU-Net. Rather, a constrained residual model improved the under-represented CC class without changing the reliable TC and WT masks. This separation of responsibilities is useful when errors are predominantly subtype confusions inside an already detected tumor.

OOF and official results also reveal a substantial domain gap. OOF CC DSC improved to 0.253, but official CC DSC was 0.144. Presence prevalence and phenotype mixture differed between development cohorts, making thresholds sensitive to case composition. ZSplit supplied interpretable signed contrasts, but component audits showed that its magnitude alone did not reliably distinguish true and false rare components. It should therefore be regarded as one contextual feature among MRI, uncertainty, anatomy, and cross-fold agreement.

The method has four limitations. First, CC and especially ED remain datalimited. Second, hard thresholds were calibrated on development data and may not transfer perfectly. Third, the adapter can only recover rare voxels already represented as NETC; it cannot repair rare tissue predicted as background or ED. Fourth, the five-fold teacher is computationally expensive. Future work should use phenotype-stratified folds, calibrated uncertainty, dedicated ED modeling, and teacher distillation while retaining the exact anatomy invariants.

## 5 Conclusion

We introduced an anatomy-preserving rare-subtype adaptation pipeline for pediatric brain tumor segmentation. Frozen nnU-Net folds provided reliable tumor anatomy and OOF error priors; component consensus and a residual adapter recovered selected ET and CC voxels. Restricting the output transition graph made TC and WT preservation mathematically explicit and deployment-safe. The approach improved OOF rare-class performance and produced a verified Docker submission, while the remaining official CC/ED gap identifies the next target for generalization.

## Acknowledgements

Data used in this publication were obtained as part of the BraTS 2026 Challenge project through Synapse ID syn74274097. We thank the BraTS organizing committee, the contributing clinical consortia, data annotators, and infrastructure teams. The author thanks ISSAI, Nazarbayev University, Astana, Kazakhstan, for providing the computing resources.

## Declaration on Generative AI

The authors used OpenAI during software development and manuscript preparation. Its uses included code assistance, manuscript organization, language refinement, and consistency checks. The authors inspected and verified the manuscript content against the implementation, experiment records, and official evaluation results. They take full responsibility for the content, scientific claims, references, and submitted software. No generative-AI system is listed as an author.

